# Rbf/E2F1 control growth and endoreplication via steroid-independent Ecdysone Receptor signalling in *Drosophila* prostate-like secondary cells

**DOI:** 10.1101/2021.03.27.437310

**Authors:** Aashika Sekar, Aaron Leiblich, Josephine E.E.U. Hellberg, Dhruv Sarma, Cláudia C. Mendes, S. Mark Wainwright, Carina Gandy, Deborah C.I. Goberdhan, Freddie C. Hamdy, Clive Wilson

## Abstract

Dysregulation of cell cycle components results in the development and progression of several cancer types. Unusually, loss of the tumour suppressor gene, *Retinoblastoma* (Rb), and consequent activation of transcription factor E2F1 have been linked to late-stage tumour progression in prostate cancer, rather than early-stage events. This change is associated with an androgen-independent form of cancer, castration-resistant prostate cancer (CRPC), which frequently still requires androgen receptor (AR) signalling. We have previously shown that binucleate secondary cells (SCs) of the *Drosophila melanogaster* male accessory gland (AG) share several functional and signalling similarities with human prostate epithelial cells. Upon mating, SC growth regulation switches from a steroid-dependent to a steroid-independent form of Ecdysone Receptor (EcR) control that induces genome endoreplication. Here, we demonstrate that the *Drosophila* Rb homologue, Rbf, and E2F1, as well as cell cycle regulators, Cyclin D (CycD) and Cyclin E (CycE), are key mediators of SC growth and endoreplication both in virgin and mated males. Importantly, we show that the CycD/Rbf/E2F1 axis requires the EcR, but not ecdysone, to trigger CycE-dependent endoreplication and associated growth in SCs after mating, mirroring changes in CRPC. We also demonstrate that excess Rbf activity reversibly suppresses binucleation in adult SCs. Overall, our work reveals mechanistic parallels between the physiological switch to hormone-independent EcR signalling in SCs, and the pathological switch seen in CRPC, and suggests that the latter may represent the dysregulation of a currently unidentified physiological process, which permits AR signalling when androgen levels are low.

## Introduction

During the early stages of prostate cancer, tumour growth requires the androgenic steroids and the androgen receptor (AR). Although androgen-deprivation therapy (ADT), a mainstay treatment for advanced prostate cancer, has an initial response rate of 90%, within two years, most cases progress to an aggressive and incurable form of prostate cancer, castration-resistant prostate cancer (CRPC) (Zong & Goldstein, 2013). Several processes play a role in the development of CRPC and in most cases, AR signalling activated in an androgen-independent manner remains essential for the maintenance and progression of the tumour (Zong & Goldstein, 2013). The molecular basis of this AR-controlled growth can involve *AR* gene amplification or increased expression, AR mutations, AR signalling pathway changes, or AR cofactors leading to androgen hypersensitivity or constitutive AR activity (Altintas et al., 2012; Zong & Goldstein, 2013).

Loss of the tumour suppressor gene *Retinoblastoma* (*Rb*) is commonly observed in several cancer types and is often considered essential for the early development of cancer (Burkhart & Sage, 2008; Sharma et al., 2010). However, in prostate cancer, *Rb* loss is associated with late-stage prostate cancer progression (Sharma et al., 2010). The role of Rb in the context of cell cycle regulation has been extensively studied (Giacinti & Giordano, 2006; Harbour & Dean, 2000a). Rb negatively regulates proteins in the E2F transcription factor family, which are involved in activating genes that are essential for the progression of the Synthesis (S)-phase of the cell cycle, including CyclinE (CycE) (Giacinti & Giordano, 2006; Harbour & Dean, 2000b). During Gap1 (G1)-phase, Rb is gradually phosphorylated, first by the growth factor-stimulated Cyclin D (CycD)/cyclin dependent kinase (cdk) 4/6 complex and then by CycE/cdk2 in late G1. It remains hyperphosphorylated until the Mitotic (M)-phase. Hyperphosphorylation of Rb changes its conformation and releases E2F factors, thereby enabling E2F-dependent transcriptional activity (Giacinti & Giordano, 2006).

In early prostate cancer, Rb tightly regulates E2F1, which in addition to CycE, also controls the expression levels of *AR*, therefore linking Rb/E2F and AR signalling. Loss of *Rb* during cancer progression leads to an unsupervised activation of E2F, thereby increasing the protein levels of AR (Sharma et al., 2010). This promotes prostate cancer cellular growth and proliferation, even after ADT, at least partly explaining the link between loss of *Rb* and CRPC (Sharma et al., 2010; Thangavel et al., 2017).

The secretory, paired male accessory gland (AG) of the fruit fly *Drosophila melanogaster* shares several functional similarities with the prostate and the seminal vesicles, the mammalian accessory glands (Leiblich et al., 2019; Ravi Ram & Wolfner, 2009; Redhai et al., 2016; Wigby et al., 2020; Wilson, Leiblich, Goberdhan, & Hamdy, 2017). The AG is formed from a monolayer epithelium consisting of two distinct octoploid binucleate cell-types: main cells (MCs) (~1000 cells/AG lobe) and secondary cells (SCs) (~40 cells/AG lobe) (Taniguchi et al., 2012). The prostate gland and the SCs, unlike the MCs, increase in size as adults age (Leiblich et al., 2012). Upon mating, the SCs grow further, partly as a result of genome endoreplication in 25% of SCs, which increases their secretory activity and therefore enhances replenishment of the AG luminal contents (Leiblich et al., 2019). The BMP pathway and the fly steroid receptor, the Ecdysone Receptor (EcR), play an essential role in regulating SC growth and endoreplication. In virgin males, the ecdysone hormone is required for growth. In mated males, however, SC growth mediated by endoreplication is ecdysone-independent (Leiblich et al., 2019), unexpectedly mirroring the transition to castration-resistant prostate cancer. Furthermore, BMP signalling, which is also implicated in CRPC, elevates EcR protein levels and promotes endoreplication in SCs (Lee et al., 2013; Leiblich et al., 2019).

Since endoreplicating cells utilise a variant of the cell cycle machinery to drive DNA replication (Edgar & Orr-Weaver, 2001; Zielke, Edgar, & DePamphilis, 2013), we investigated the molecular mechanisms controlling ecdysone-independent, EcR-mediated endoreplication in SCs. Here we demonstrate that CycD, Rbf and E2F1 are essential for SC endoreplication and growth after mating, with Rbf playing an additional role in controlling binucleation. Although CycE appears to function downstream of EcR, CycD/Rbf/E2F1 require EcR to control endoreplication; increased E2F1 expression elevates EcR protein levels and in turn, EcR promotes hormone-independent activation of CycE-mediated endoreplication. Our data therefore reveal a physiological mechanism involving Rbf/E2F1-activated control of hormone-independent EcR-mediated endoreplication and growth, which mirrors pathological growth-promoting changes in AR signalling associated with Rb loss in CRPC.

## Results

### EcR signalling requires CycE to induce SC growth and endoreplication

CycE has been implicated in the regulation of endoreplication in several *Drosophila* cell types, including SCs (Edgar, Zielke, & Gutierrez, 2014; Leiblich et al., 2019). We further studied the role of CycE in the control of SC growth and endoreplication using the esg^ts^F/O driver to overexpress transgenes specifically in adult SCs under the control of the yeast GAL4 transcription factor (Leiblich et al., 2019, 2012). This line ubiquitously expresses a temperature-sensitive form of the GAL4 inhibitor GAL80, which blocks *esg*-GAL4-dependent transgene expression until the temperature is shifted to 28.5°C, which inhibits GAL80 function. Newly eclosed males were switched from 18°C to 28.5°C for 6 days to express the transgene in SCs. SC growth was measured as a ratio of SC nuclear growth to adjacent MC nuclear growth (Leiblich et al., 2019).

Using this system, we confirmed that *CycE* knockdown resulted in a decrease in SC nuclear growth in both virgin and mated males (Fig.S1A-D, E). Furthermore, consistent with previous observations (Leiblich et al., 2019), *CycE*-RNAi expression completely inhibited mating-dependent endoreplication normally seen in 25% of SCs (Fig.S1A-D, F), as assayed by nuclear incorporation of 5-ethynyl-2′-deoxyuridine (EdU), a synthetic analogue of thymidine delivered in the food. Contrary to our previous findings, SC growth in knockdown cells was significantly increased after mating, although this change was only detectable because of the reduced number of genotypes being compared. CycE is therefore required for SC nuclear growth in virgin and mated males, and for mating-dependent endoreplication, but some endoreplication-independent growth in mated males does not appear to be mediated by CycE.

Since we had previously shown that the EcR is necessary and sufficient to induce endoreplication in SCs, we determined whether CycE is dependent on the EcR to promote SC endoreplication. When *CycE* was co-overexpressed with *EcR*-RNAi, 100% of SCs endoreplicated (Fig.1A-D, F), and CycE-induced nuclear overgrowth was not inhibited (Fig.1A-D, E), demonstrating that CycE acts downstream or independently of EcR in this growth regulatory pathway.

**Figure 1:**
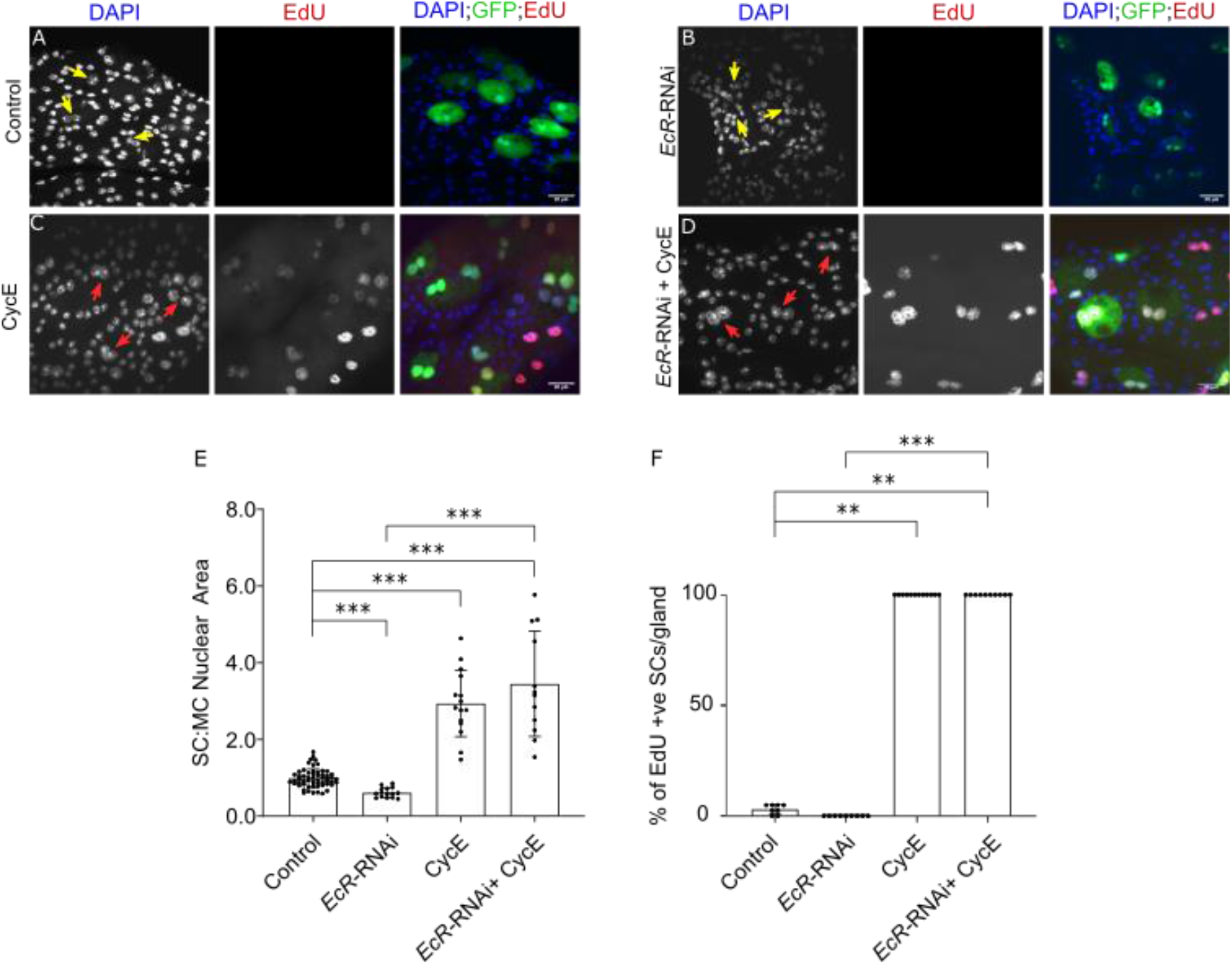
EcR is not required for CycE-induced SC growth and endoreplication. (A-D) Images show distal tip of AGs from 6-day old adult virgin males expressing nuclear GFP alone (control; this also stains the cytosol) or in combination with additional transgenes in SCs under the control of the esg^ts^F/O driver and stained for EdU incorporation in SCs. Nuclei are stained with DAPI (blue). Red arrows point to EdU^+^ SC nuclei and yellow arrows point to EdU^−^ SC nuclei for each transgene. (E-F) Histograms depicting the mean ratio of the size of SC nuclei relative to neighbouring MC (E) and the mean % of EdU^+^ SCs per gland (F) of virgin males expressing no transgene, *EcR*-RNAi, CycE or *EcR*-RNAi + CycE in SCs. Knocking down *EcR* does not affect CycE-mediated SC growth and endoreplication. Welch ANOVA; Games-Howell post hoc test; n≥15 cells (E). Kruskal Wallis test; Dunn’s post hoc test; n≥8 glands (F). Scale bars correspond to 20 μm. The error bars show the standard deviation within the sample. 0.0001<***p≤0.001; ****p≤0.0001.

### Activated Rbf suppresses binucleation in SCs

To test the effect of upstream CycE regulators on SC growth and endoreplication, we initially expressed under GAL4/UAS control the constitutively activated Rbf^CA^ protein, which has three of its four phosphorylation sites substituted, thereby making it refractory to regulation by CycE/Cdk2 and CycD/Cdk4 (Xin, Weng, Xu, & Du, 2002). Interestingly, this resulted in 63± 12% of SCs becoming mononucleated after 6 days of adult expression (Fig.2A,C,C’,I). Remarkably, we observed that this mononucleate phenotype could be partially reversed to a binucleate state, when the expression of Rbf^CA^ was blocked again (Fig.2E,F,K). To test whether inhibition of E2F1 is involved in this phenomenon, it was knocked down in SCs, resulting in 12 ± 3% of SCs becoming mononucleated (Fig.2H,I). Since mononucleation is never observed in controls, this suggests that E2F1 inhibition plays a role in Rbf-induced mononucleation, and E2F1 is involved in normal maintenance of binucleation in SCs.

**Figure 2:**
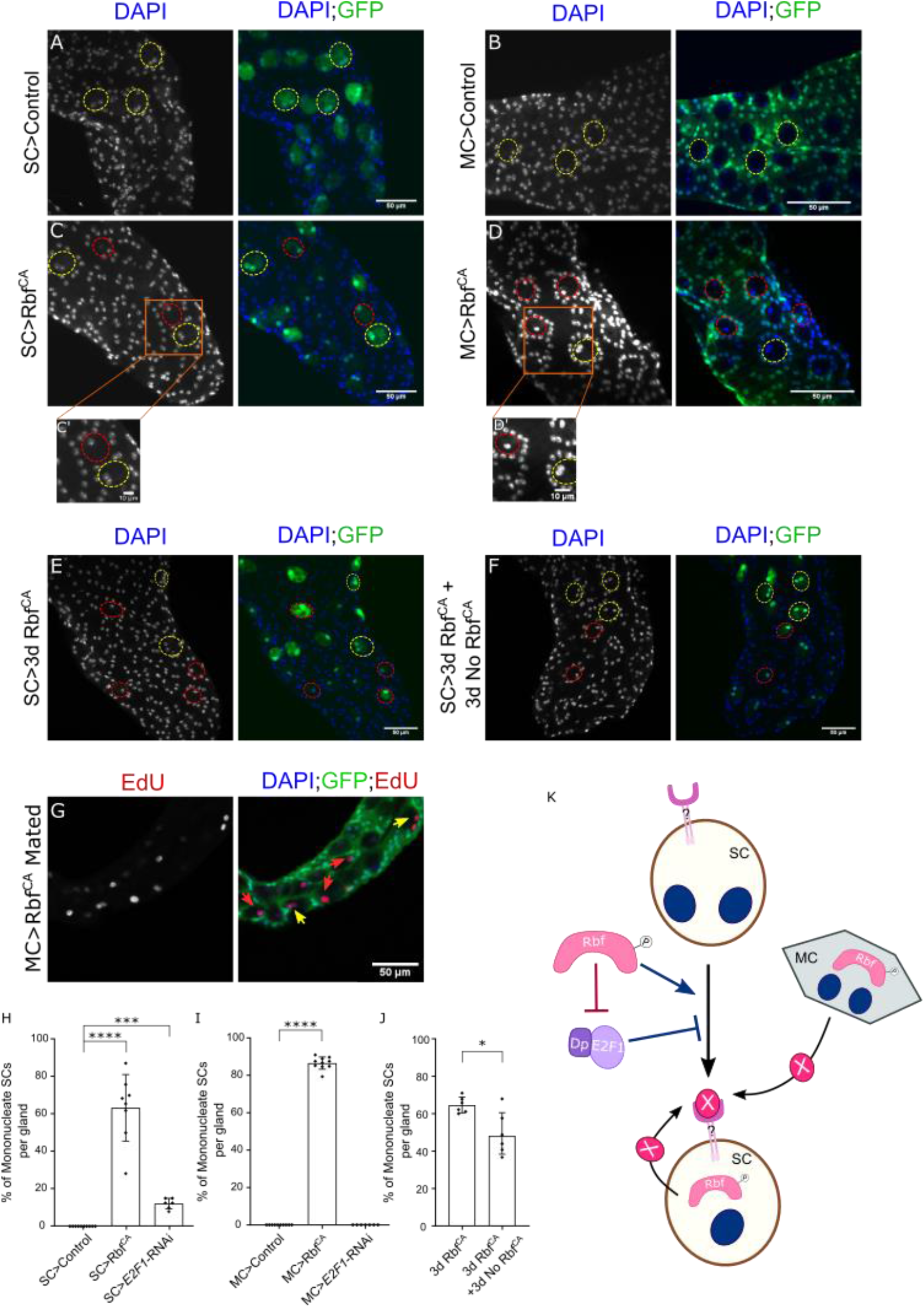
Rbf/E2F1 axis maintains adult SC nucleation state. (A-D) Images show distal tip of AGs from 6-day-old adult virgin males of control glands or glands expressing Rbf^CA^ either under the control of the esg^ts^F/O driver (A,C) or the Acp26Aa-GAL4 driver (B,D) which activates nuclear GFP production (stains cytoplasm in A,C). Orange boxes outline the area zoomed in the inset (C’,D’). (E) Image shows distal tip of an AG from a 3-day-old adult virgin male expressing Rbf^CA^ in SCs under the esg^ts^F/O control. (F) Image shows distal tip of an AG from a 6-day-old virgin male expressing Rbf^CA^ in SCs for 3 days since eclosion, followed by 3 days of no Rbf^CA^ expression. (G) Image shows the distal tip of an AG of a 6-day-old multiply-mated male expressing Rbf^CA^ under the control of the Acp26Aa-GAL4 driver. The AG was stained for EdU, revealing that mononucleate SCs are able to endoreplicate. (A-G) Nuclei are stained with DAPI (blue). Dashed red ellipses mark the outline of mononucleate SCs; dashed yellow ellipses mark the outline of binucleate SCs; red arrows mark the nuclei of mononucleate SCs; yellow arrows mark the nuclei of binucleate SCs. (H-J) Histogram depicting the mean % of mononucleate SCs per gland of virgin males of control glands or glands expressing transgenes either in SCs (H-I) or MCs (J). Expression of Rbf^CA^ either in SCs or MCs induces SC mononucleation in a reversible manner, but no such effects are observed in MCs. Knocking down *E2F1* in SCs promotes SC mononucleation but no effect is observed when knocked down in MCs. Kruskal Wallis test; Dunn’s post hoc test; n≥6 glands (H-J). Scale bars correspond to 50 μm (A-G) and 10 μm (C’, D’). 0.01≤*p≤0.05; ****p≤0.0001. (K) Schematic detailing nucleation state regulation in adult SCs. Expression of a constitutively active form of Rbf, Rbf^CA^, which has mutations in three of its four CDK phosphorylation sites, in SCs is able to induce mononucleation. This phenotype is negatively regulated by E2F1. Additionally, expression of Rbf^CA^ in MCs is able to activate mononucleation in SCs, presumably by secreting ‘X’ which could activate an unknown receptor (?) which regulates SC nucleation state. It is possible that Rbf^CA^ when expressed in SCs also promotes mononucleation via this unknown receptor.

In order to determine whether the nucleation state of MCs is also affected by Rbf, Rbf^CA^ was overexpressed in these cells, using a GAL80^ts^-regulated *Acp26Aa*-GAL4 MC driver (Fig.2B,D,D’,J) (Chapman et al., 2003; Corrigan et al., 2014). Surprisingly, there was no effect on MC binucleation, but 86 ± 3% SCs became mononucleated. These SCs could still endoreplicate upon mating (Fig.2G), demonstrating that adult endoreplication is not affected by the nucleation state of SCs and that the mononucleation phenotype is unlikely to be mediated by transfer of endoreplication-inhibiting Rbf^CA^ to SCs. Expression of *E2F1*-RNAi in MCs did not induce mononucleation either in SCs or MCs (Fig.2 J; Fig.S2C). In summary, these results suggest that Rbf can control SC nucleation state in two different ways; cell-autonomous regulation in which E2F1 appears to be partly involved, and paracrine regulation, which may not require E2F1 inhibition, and where the downstream signals are unknown.

### CycD, Rbf and E2F1 are essential for the regulation of SC growth and endoreplication

Having observed a role for *Rbf* and *E2F1* in SC binucleation, we tested the role of these genes and their upstream cell cycle regulator CycD in SC growth and endoreplication. Knocking-down *CycD* in SCs inhibited SC nuclear growth and abolished mating-dependent endoreplication (Fig.3H, K; Fig.S1J). However, similar to when *CycE* was knocked down, we observed that expression of *CycD*-RNAi did not fully suppress all mating-dependent SC growth. To test the effect of increased CycD activity, CycD was overexpressed with its kinase partner, Cdk4, in SCs (Fig.3G, H). All the cells endoreplicated extensively, resulting in polytene-like chromosomes in both virgin and mated males (Fig.3G’,K; Fig.S1I). Taking these results together, CycD is necessary and sufficient to induce SC growth and endoreplication.

**Figure 3:**
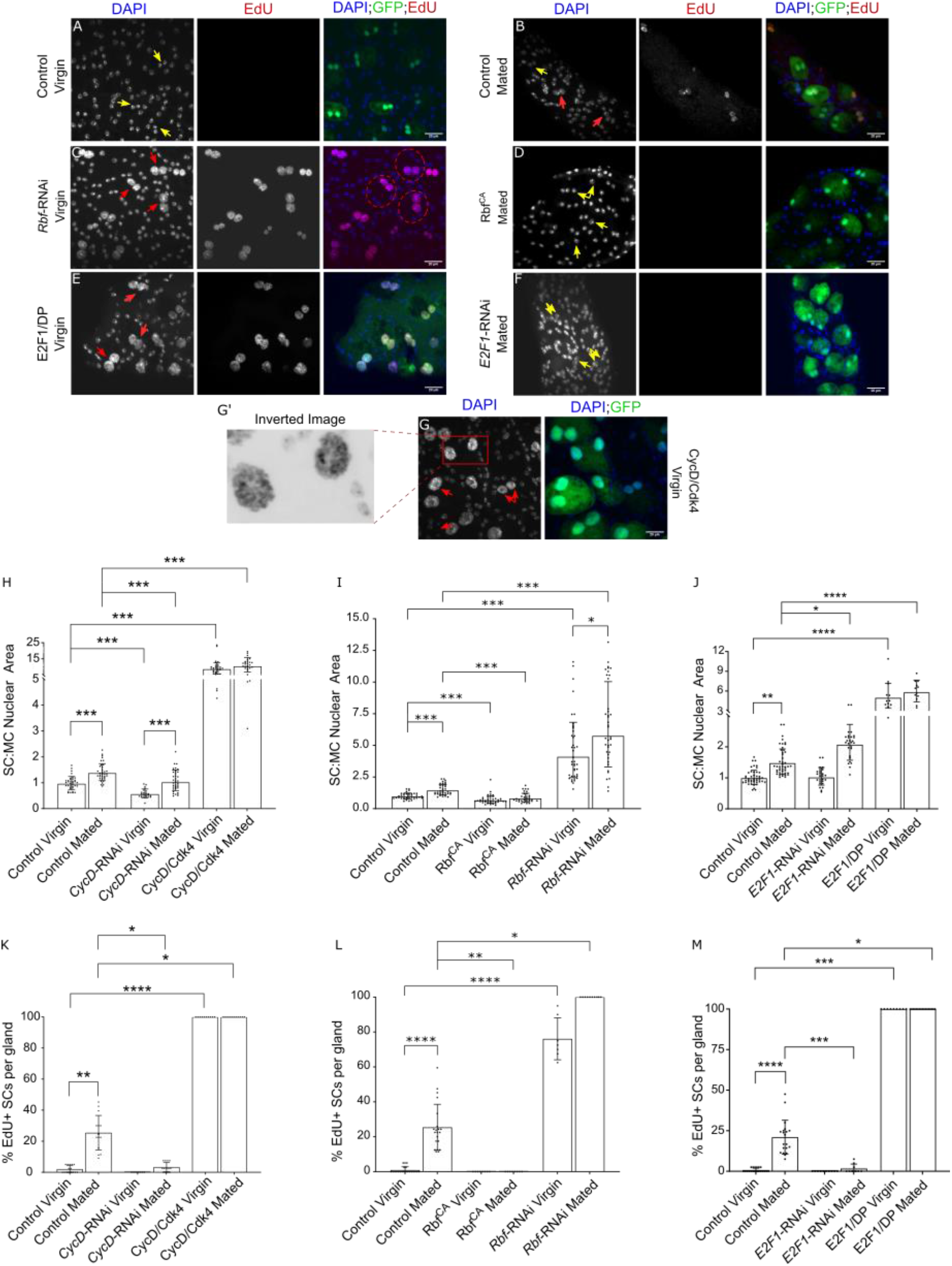
Cell cycle components regulate SC growth and endoreplication. (A-G) Images show distal tip of AGs from 6-day old adult virgin males expressing nuclear GFP alone (control; this also stains the cytosol) or in combination with additional transgenes in SCs under the control of the esg^ts^F/O driver and stained for EdU incorporation in SCs. Nuclei are stained with DAPI (blue). Red arrows point to EdU^+^ SC nuclei and yellow arrows point to EdU^−^ SC nuclei for each transgene (G’) Magnified negative image of orange square in image G to highlight the polytene-like nuclei in SCs overexpressing CycD/cdk4. (H-M) Histograms depicting the geometric mean ratio of the size of SC nuclei relative to neighbouring MC nuclei (H-J) and mean % of EdU+ SCs per gland (K-M) of virgin and mated males of control glands and glands expressing different transgenes in SCs. CycD is necessary and sufficient to promote SC growth and endoreplication. Knocking down *CycD* is unable to completely suppress SC growth between the respective virgin and mated males, which as seen in K, is associated with mating-induced, endoreplication-independent SC growth. Rbf negatively regulates SC growth and endoreplication in virgin and mated males. E2F1 is not necessary for SC growth in virgins, but in mated males, it appears that overall it negatively regulates growth, which EdU analysis indicates is endoreplication-independent. Welch ANOVA on log-transformed data; Games-Howell post hoc test (H-J). Kruskal Wallis test; Dunn’s post hoc test (K-M). n≥24 cells (H); n≥33 cells (I); n≥12 cells (J); n≥9 glands (K); n≥7 glands (L); n≥9 glands (M). Scale bars correspond to 20 μm. The error bars show the geometric standard deviation (H-J) and standard deviation (K-M) within the sample. 0.01<*p<0.05; 0.001<**p≤0.01; 0.0001<***p≤0.001; ****p≤0.0001.

Overexpression of Rbf^CA^ suppressed mating-dependent endoreplication and all SC nuclear growth in virgin and mated males (Fig.3 D, I, L). In these experiments, the ratio of the size of the single nucleus in mononucleate SCs to the total nuclear area of both nuclei in the adjacent MC was measured, since the reduction observed in mononucleate cells using this modification to the measurement of nuclear growth correlated with the reduction in cell area found in mononucleate versus binucleate SCs (Fig.S3). Expressing *Rbf*-RNAi resulted in an increase in growth and endoreplication in all SCs in both virgin and mated males (Fig.3C,I,L). Hence, Rbf normally limits SC endoreplication and growth, and its hyperactivation suppresses all forms of SC growth in both virgins and mated males.

To increase E2F1 activity in SCs, it was co-expressed with its dimerization partner, Dumpy (DP). Increased SC growth was observed and all SCs underwent endoreplication in both virgin and mated males (Fig.3 E, J, M). Surprisingly, *E2F1* knockdown had no effect on SC growth in virgin males and growth was increased in mated males versus mated controls. However, all endoreplication was suppressed after mating (Fig.3 F, J, M). Therefore, E2F1 is not necessary for endoreplication-independent SC growth in virgin and mated males. Indeed, since knockdown cells in mated males have larger nuclei than controls after mating, E2F1 appears to normally suppress growth that occurs without endoreplication after mating, as well as driving endoreplication-associated growth. Knockdown of the other Rbf-regulated *E2F* gene in *Drosophila*, *E2F2*, had no effect on SC growth and endoreplication (Fig.S4), suggesting that this gene is not involved in these processes. Since *E2F1* knockdown cells in mated males have larger nuclei than cells in mated controls, it appears that E2F1 normally suppresses growth that occurs without endoreplication after mating, as well as driving endoreplication-associated growth.

We conclude that the cell cycle components CycD, Rbf and E2F1 all play important roles in the regulation of adult SC growth and endoreplication, although CycD/Rbf do not seem to act via E2F1 to control endoreplication-independent SC growth in virgin and mated males.

### BMP signalling is downstream of CycD, but upstream of E2F1 in the regulation of EcR expression, growth and endoreplication in SCs

Since BMP signalling also drives growth and endoreplication in SCs (Leiblich et al., 2019), we investigated whether the BMP signalling pathway interacts with the CycD/Rbf/E2F1 signalling axis in these cells. When we co-expressed E2F1/DP with a BMP antagonist, Daughters against Decapentaplegic (Dad) in SCs, we observed that nuclear growth was much higher than the controls and not significantly decreased from cells expressing E2F1/DP alone (Fig.4B-D, 4I). Furthermore, all the SCs expressing E2F1/DP and Dad endoreplicated in virgin males (Fig.4B-D, 4L).

**Figure 4:**
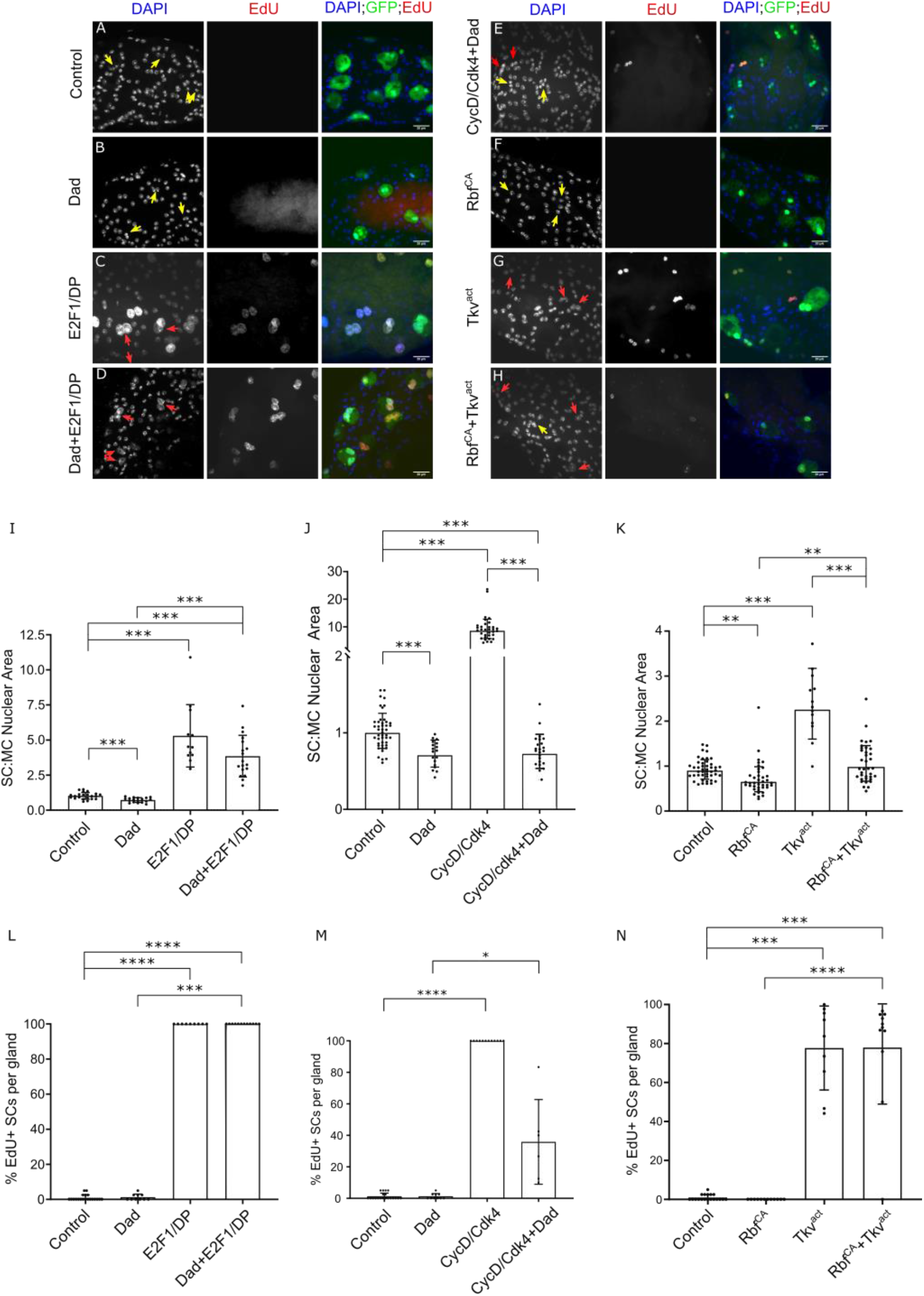
Interaction between CycD/Rbf/E2F1 and BMP signalling in the regulation of SC growth and endoreplication. (A-H) Images show distal tip of AGs from 6-day old adult virgin males expressing nuclear GFP alone (control; this also stains the cytosol) or in combination with additional transgenes in SCs under the control of the esg^ts^F/O driver and stained for EdU incorporation in SCs. Nuclei are stained with DAPI (blue). Red arrows point to EdU^+^ SC nuclei and yellow arrows point to EdU^−^ SC nuclei for each transgene. (I-N) Histograms depicting the mean SC:MC nuclear area (I), geometric mean SC:MC nuclear area (J-K) and mean % of EdU+ SCs per gland (L-N) of virgin males of control glands and glands expressing different transgenes in SCs. Dad is not able to suppress E2F1/DP-mediated SC growth and endoreplication, but Dad is able to suppresse CycD/cdk4-mediated SC growth and endoreplication. Rbf^CA^ suppresses most of the growth induced by Tkv^act^, however Rbf^CA^ is not able to suppress Tkv^act^-induced endoreplication. Welch ANOVA; Games-Howell post hoc test (I). Welch ANOVA on log-transformed data; Games-Howell post hoc test (J-K). Kruskal Wallis test; Dunn’s post hoc test (L-N). n≥12 cells (I); n≥18 cells (J); n≥13 cells (K); n≥9 glands (L); n≥9 glands (M); n≥6 glands (N). Scale bars correspond to 20 μm. The error bars show the standard deviation (I, L-N) and geometric standard deviation (J-K) within the sample. 0.01<*p<0.05; 0.001<**p≤0.01; 0.0001<***p≤0.001; ****p≤0.0001.

By contrast, when CycD/Cdk4 was co-expressed with Dad, Dad completely suppressed all SC nuclear growth induced by CycD/Cdk4 alone (Fig.4E, J; Fig.S1I). Additionally, only 20% of SCs showed any endoreplication (Fig.4E, M; Fig.S1I), despite the fact that CycD/Cdk4 can induce multiple rounds of endoreplication in the absence of Dad (Fig.3G’). We attribute this low level of DNA synthesis to Dad expression incompletely suppressing BMP signalling in some SCs during the six-day expression period. Therefore, we conclude that CycD acts upstream of BMP signalling’s effects on nuclear growth and endoreplication; in turn, BMP signalling is upstream of E2F1/DP.

In order to further investigate how the BMP pathway interacts with the CycD/Rbf/E2F1 axis, we co-expressed Rbf^CA^ and a constitutively active form of a BMP Type-I receptor, Thickveins, (Tkv^act^) in SCs. This resulted in nuclear growth similar to controls in virgin males (Fig.4F-H, K). This growth was much reduced compared to when only Tkv^act^ was expressed; indeed, it was not significantly different from the growth observed when Rbf^CA^ was expressed alone. However, surprisingly 80% of the SCs still endoreplicated, phenocopying the effect of Tkv^act^ alone. (Fig.4F-H, N). Overall, we conclude that even though BMP signalling has a dominant effect on endoreplication when co-modulated with CycD/Rbf signalling, Rbf^CA^ overexpression suppresses the growth-promoting effects of BMP signalling on SC nuclei independently of endoreplication, suggesting that these two signals interact in different ways to control these two cellular processes.

Elevated BMP signalling is known to dramatically increase EcR protein expression in SCs presumably contributing to the effects of this signalling pathway on SC growth and endoreplication (Leiblich et al., 2019). Since E2F1 functions downstream of BMP signalling in endoreplication control, we hypothesised that Rbf/E2F1 might also affect EcR protein levels. Indeed, EcR protein, detected using a pan-EcR antibody, was greatly increased in comparison to control cells, when either E2F1/DP or *Rbf*-RNAi were overexpressed in SCs, with EcR present in the cytoplasm, as well as in all nuclei (Fig.5A-C). Expressing Rbf^CA^ in SCs completely suppressed EcR protein expression, mirroring the phenotype observed with expression of *EcR*-RNAi (Fig.5D-E). However, *E2F1*-RNAi did not have any obvious effect on EcR protein levels in SCs (Fig.5F), perhaps explaining the weaker effects of this knockdown on SC nuclear growth compared to Rbf^CA^ (Fig.3I, J). To confirm specific staining with the EcR antibody, we expressed E2F1/DP with *EcR*-RNAi in SCs; no EcR expression was observed (Fig.5G). We conclude that the BMP-regulated CycD/Rbf/E2F1 axis controls not only SC nuclear growth and endoreplication, but also expression of EcR protein, another key player in SC growth.

**Figure 5:**
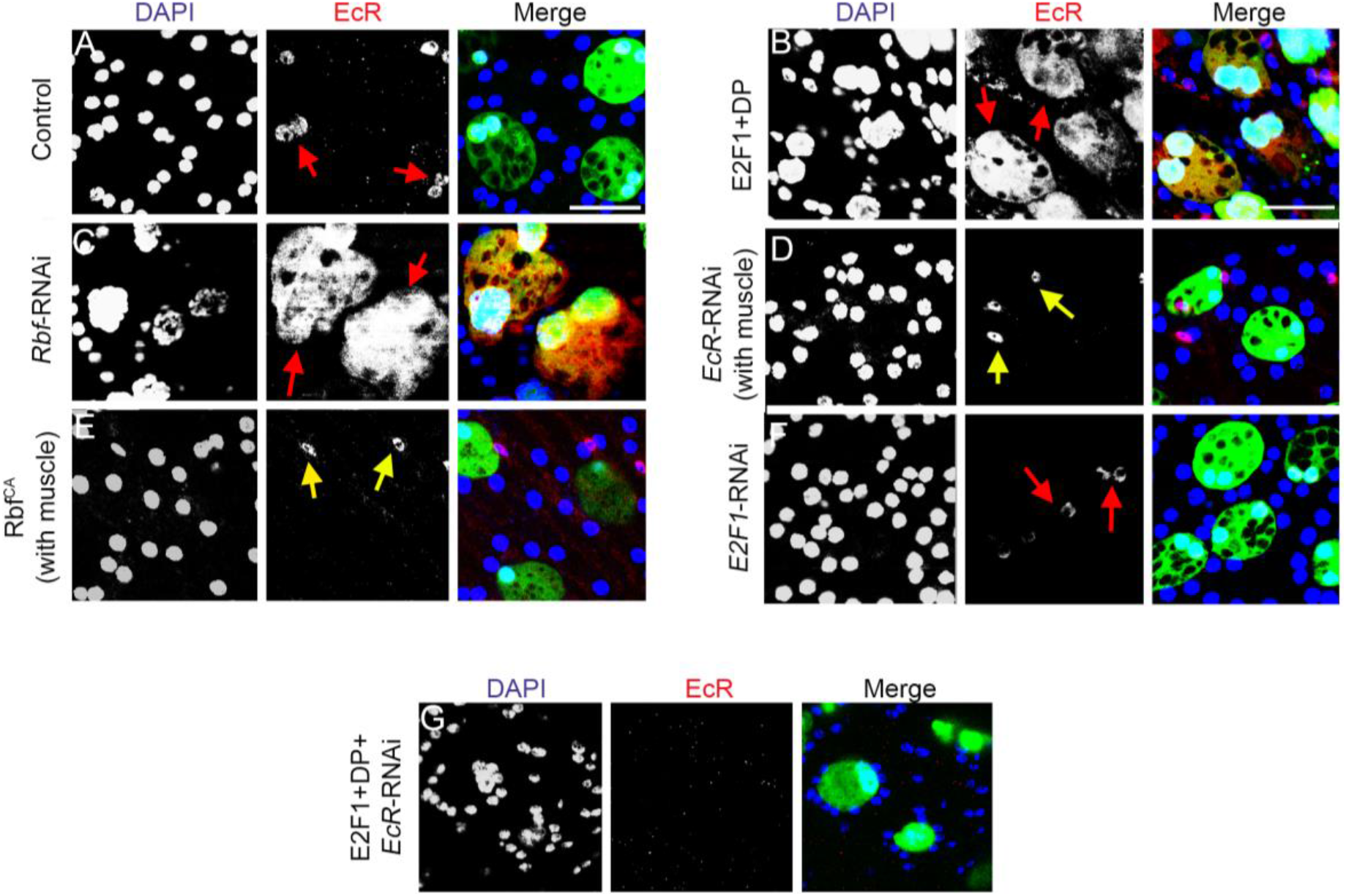
Rbf/E2F1 regulates EcR protein levels in SCs. (A-G) Images show distal tip of AGs from 6-day old adult virgin males expressing nuclear GFP alone (control; this also stains the cytosol) or in combination with additional transgenes in SCs under the control of the esg^ts^F/O driver and stained with a pan-EcR antibody (red) in SCs. Nuclei are stained with DAPI (blue). Red arrows mark SCs expressing EcR. Yellow arrows mark muscle cells expressing EcR as a control for EcR staining in glands which do not have EcR expression in SCs (E,F). Rbf negatively regulates EcR protein levels in SCs (C,F). Although overexpression of E2F1/DP results in an increase in EcR expression in SCs, expression of *E2F1*-RNAi does not appear to affect EcR protein levels. Scale bars correspond to 50 μm.

### Rbf and E2F1 function upstream of the EcR in the hormone-independent regulation of SC endoreplication

As discussed earlier, *Rb* loss and E2F1 activation increase AR levels in prostate cancer and are associated with CRPC, in which the AR can promote cell growth and proliferation in a hormone-independent fashion. Hence, we tested the functional interactions between the CycD/Rbf/E2F1 signalling axis and the EcR steroid receptor to determine whether these might be involved in ecdysone-independent growth and endoreplication in SCs, mirroring the effects in CRPC.

When E2F1/DP was co-expressed with *EcR*-RNAi, SC nuclei grew to a greater size than controls or SCs expressing *EcR*-RNAi, but growth was suppressed compared to cells expressing E2F1/DP alone (Fig.6B-D,J). However, *EcR*-RNAi completely suppressed E2F1/DP-induced endoreplication (Fig.6B-D, M). Hence, we conclude that E2F1/DP can elicit some SC growth when EcR expression is suppressed, but, somewhat surprisingly, for SC endoreplication, E2F1/DP requires the EcR to drive this CycE-dependent event. Co-expressing Rbf^CA^ with EcR-C produced more variable and intermediate phenotypes, probably because each SC expresses different relative levels of these transgenes. Whereas 100% of SCs expressing EcR-C alone endoreplicated in virgin males, only 60% endoreplicated when Rbf^CA^ was co-expressed, with some glands exhibiting endoreplication in 100% of SCs and others in only 2.5%. No endoreplication was observed in SCs expressing Rbf^CA^ alone (Fig.6E-G, K, N). These results suggest that EcR-C is able to induce endoreplication even when SCs express Rbf^CA^, consistent with our finding that EcR acts downstream of E2F1 in endoreplication control. Furthermore, nuclear growth levels in SCs of these males were similar to controls, an intermediate phenotype that suggests Rbf can act somewhat independently of EcR to control growth, as we also found for E2F1 (Fig.6 J).

**Figure 6:**
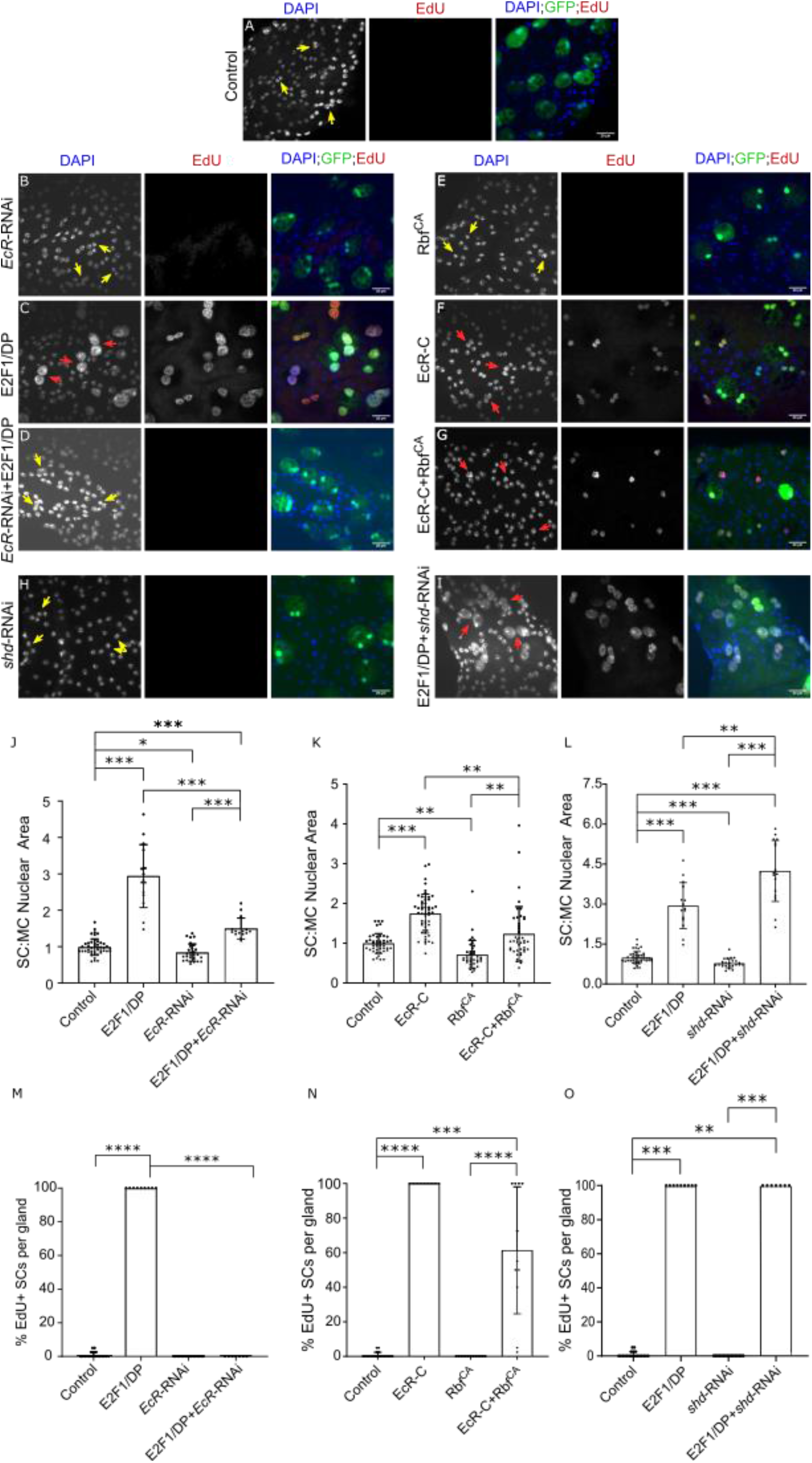
Rbf/E2F1 interacts with the EcR to regulate SC growth and hormone-independent endoreplication. (A-I) Images show distal tip of AGs from 6-day old adult virgin males expressing nuclear GFP alone (control; this also stains the cytosol) or in combination with additional transgenes in SCs under the control of the esg^ts^F/O driver and stained for EdU incorporation in SCs. Nuclei are stained with DAPI (blue). Red arrows point to EdU^+^ SC nuclei and yellow arrows point to EdU^−^ SC nuclei for each transgene. (J-O) Histograms depicting the geometric mean SC:MC nuclear area (J-K), mean SC:MC nuclear area (L) and mean % of EdU+ SCs per gland (M-O) of virgin males of control glands and glands expressing different transgenes in SCs. E2F1 can promote some growth in SCs independently of EcR. However, E2F1-mediated endoreplication in SCs requires the ecdysone-independent EcR signalling. Rbf can either interact with EcR or act independently to negatively regulate SC growth. Welch ANOVA on log-transformed data; Games-Howell post hoc test (J-K). Welch ANOVA; Games-Howell post hoc test (L). Kruskal Wallis test; Dunn’s post hoc test (M-O). n≥15 cells (J); n≥39 cells (K); n≥15 cells (L); n≥7 glands (M); n≥11 glands (N); n≥7 glands (O). Scale bars correspond to 20 μm. The error bars show the geometric standard deviation (J-K) and standard deviation (K-M) within the sample. 0.01<*p<0.05; 0.001<**p≤0.01; 0.0001<***p≤0.001; ****p≤0.0001.

Finally, since endoreplication is regulated by the EcR in a hormone-independent fashion, we hypothesised that E2F1 activation might also stimulate EcR-mediated genome replication in the absence of hormone, mirroring events in CRPC. Co-expression of E2F1/DP with an RNAi targeting *shd*, the gene encoding ecdysone 20-hydroxylase required for the last step in 20-hydroxyecdysone synthesis, had no inhibitory effect on SC endoreplication and in fact, resulted in an increase in nuclear growth compared to E2F1/DP expression alone (Fig.7), in sharp contrast to E2F1/DP co-expression with EcR-RNAi (Fig.6 D, J, M). Therefore, E2F1/DP-induced SC endoreplication is mediated by hormone-independent EcR signalling and ecdysone may, in fact, interfere with some aspects of E2F1/DP-mediated nuclear growth.

**Figure 7:**
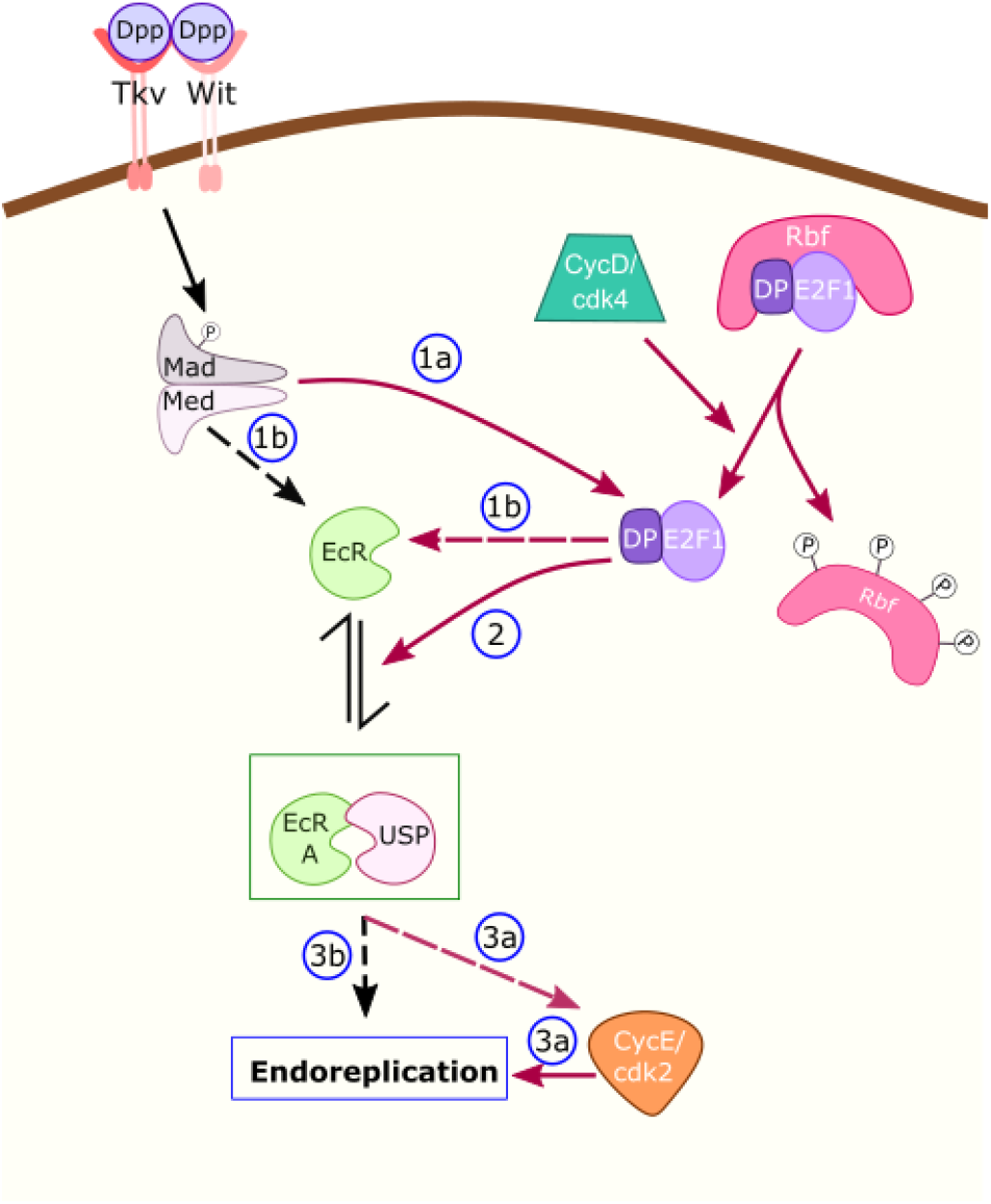
Schematic detailing cell cycle regulation of SC growth and genome endoreplication. In virgin males, CycD, Rbf and CycE regulate SC growth, but E2F1 does not play a role in this regulation. In mated males, CycD/Rbf/E2F1/CycE regulates SC growth that occurs through endoreplication. Interestingly, it appears that E2F1 requires EcR to activate endoreplication, but whether either E2F1 or EcR can exert their control through CycE to regulate endoreplication is not known. Rbf can inhibit SC growth that occurs without endoreplication in mated males, but it appears that knocking down CycD or CycE is not able to suppress this growth. Furthermore, CycD/Rbf /E2F1 is an upstream regulator of EcR, while CycE acts downstream of EcR. BMP signalling interacts with the cell cycle regulatory axis downstream of Rbf but upstream of E2F1. Therefore, in SCs, the cell cycle machinery has a distinct role from its cell cycle regulation that involves an important input from the EcR. cdk4/2- cyclin dependent kinase 4/2; CycD- Cyclin D; CycE- Cyclin E; Dp- Dumpy; Dpp-Decapentaplegic; Ec-Ecdysone; EcR-Ec receptor; Med-Medea; P- Phosphate group; Rbf-Retinoblastoma; Tkv- Thickveins; Usp-Ultraspiracle; Wit-Wishful thinking; blue ?-unknown EcR partners. Red arrows depict the pathways that regulate SC growth and endoreplication and the dotted arrows show probable signalling interactions.

## Discussion

Multiple mechanisms can contribute to the inevitable emergence of CRPC following androgen-deprivation therapy, including the late-stage loss of *Rb*, a change that occurs at much earlier stages in most other cancers. In the prostate, this genetic change increases hormone-independent AR signalling, in part by increasing AR expression levels, although other mechanisms are likely to be involved (Sharma et al., 2010). Using the *Drosophila* SC, which shares several cell biological similarities with prostate epithelial cells, we demonstrate here that the *Drosophila Rb* homologue, *Rbf*, regulates the receptor for the steroid ecdysone to suppress SC growth, including the ecdysone-indpendent growth which is induced after male flies mate (Fig.8). Most notably, Rbf inhibits hormone-independent endoreplication observed after mating, and when the CycD/Rbf/E2F1 axis is activated, the EcR is essential for the induction of endoreplication. This physiological mechanism is not observed in other *Drosophila* cell types, but does mirror changes seen in CRPC, suggesting the possibility that the latter pathological change might reflect a physiological process in human prostate epithelium that is yet to be characterised.

**Figure 8:**
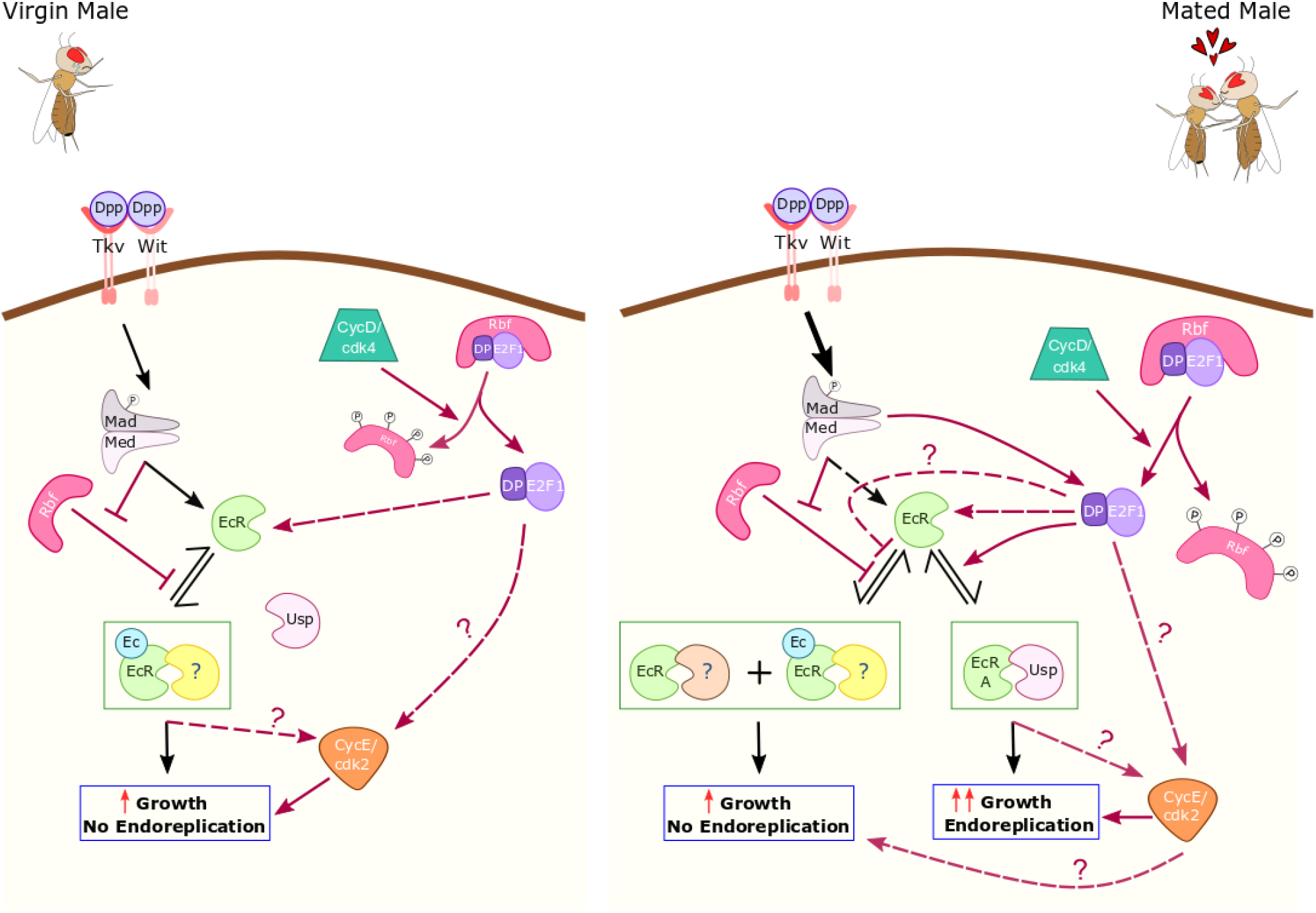
Schematic detailing nucleation state regulation in adult SCs. Expression of a constitutively active form of Rbf, Rbf^CA^, which has mutations in three of its four CDK phosphorylation sites, in SCs is able to induce mononucleation. This phenotype is negatively regulated by E2F1 and positively regulated by p53. Additionally, expression of Rbf^CA^ in MCs is able to activate mononucleation in SCs, presumably by secreting ‘X’ which could activate an unknown receptor (?) which regulates SC nucleation state. It is possible that Rbf^CA^ when expressed in SCs also promotes mononucleation via this unknown receptor.

### Activated Rbf reverses the binucleate state in adult SCs independently of its effects on growth and endoreplication

Our study has revealed an interesting additional role for Rbf in controlling the nucleation state in binucleate adult SCs. Rbf^CA^ reversibly induces almost all SCs to become mononucleate, but has no effect on MC nucleation state. Furthermore, knocking down E2F1 in SCs results in just a few SCs becoming mononucleate, whereas no effect was observed when E2F1 was knocked down in MCs. Therefore, SCs appear to require some signalling by the CycD/Rbf/E2F1 axis to retain their binucleate state in adults.

Perhaps most surprisingly, expression of the Rbf^CA^ construct in MCs results in SC mononucleation, while MCs retain their binucleate nature. In this case, those SCs, which become mononucleate, are still able to endoreplicate upon mating. Hence, it appears that an unknown secreted MC signal, which is not transferred Rbf, can direct mononucleation in SCs. Since MCs do not become mononucleated, MCs either must lack the proper machinery to process this mononucleation signal or regulate binucleation in a different way. *E2F1*-RNAi expression in MCs does not have the capacity to elicit the mononucleation signal to SCs, indicating that E2F1 is not involved in this signalling mechanism, although this might also be accounted for by insufficient knockdown of *E2F1* in MCs. Interestingly, a recent study has shown that although Rb/E2F does not initiate binucleation in mammalian cardiomyocytes, the binucleate and mononucleate cells have distinct Rb/E2F-mediated transcriptional programmes (Windmueller et al., 2020), thereby emphasising the importance of this pathway in different nucleation states.

### CycD/Rbf/E2F1 signalling axis and CycE regulate transition from hormone-dependent to hormone-independent EcR signalling in SCs

We have demonstrated that CycD and CycE, are necessary and sufficient for the induction of endoreplication in SCs, and that they are also required for ecdysone-dependent SC nuclear growth in virgin males, even though no detectable DNA replication occurs in these cells (Fig.8). However, they are not needed for at least some of the endoreplication-independent SC growth that occurs in mated males. In breast cancer and prostate cancer, CycD1 and CycE have been demonstrated to regulate both ligand-dependent and ligand-independent steroid receptor signalling, the latter playing a vital role in the initiation of hormone-refractory form of cancer growth (Bindels, Lallemand, Balkenende, Verwoerd, & Michalides, 2002; Foster, Henley, Ahamed, & Wimalasena, 2001; Petre, Wetherill, Danielsen, & Knudsen, 2002).

CycD-regulated Rbf is also involved in normally suppressing these SC growth regulatory functions, although expression of a constitutively active form of Rbf appears to be able to block all growth, even in mated males. By contrast, although E2F1 is necessary and sufficient to promote endoreplication in SCs, it is not essential for growth that occurs in virgins but appears to inhibit endoreplication-independent growth in mated males. This suggests that Rbf and E2F1 can act independently to fulfil some of their activity. Rb functions that occur independently of its E2F transcriptional activity control have been observed in Rb-mediated cell cycle arrest and chromatin stability in mammalian cells (Dick & Rubin, 2013). The E2F1-independent Rbf activity that regulates chromatin condensation appears to be conserved in *Drosophila* as well (Longworth, Herr, Ji, & Dyson, 2008). There is also some evidence pertaining to E2F1 activity independent of Rbf during early embryogenesis (Shibutani, Swanhart, & Duronio, 2007). However, canonical Rbf/E2F1 signalling is also active in SCs and is required for endoreplication-dependent SC growth in mated males.

Rbf and E2F1 can mediate the effects on SC growth and endoreplication partly by regulating the EcR protein levels, as previously observed with the BMP signalling (Leiblich et al., 2019). Epistasis experiments suggest that the BMP and EcR pathways interact with the CycD/Rbf/E2F1/CycE axis in SCs. BMP signalling intersects the cell cycle regulatory pathway upstream of E2F1 activity and CycE, and downstream of Rbf and CycD. EcR lies downstream of CycD/Rbf/E2F1, but upstream of CycE in the regulation of endoreplication and growth in SCs. Furthermore, we confirmed that E2F1 induces endoreplication and growth in an ecdysone-independent fashion, which mirrors the effects observed with loss of Rb in CRPC, which induces unsupervised activation of E2F activity.

Comprehensive genomic profiling of prostate cancer tumours, cell lines and xenografts has revealed that Rb signalling is functionally altered in 34% in primary tumours and in 74% of castration-resistant tumours, thereby stressing the importance of Rb in the emergence of CRPC (Taylor et al., 2010). Studies in prostate cancer cell lines have demonstrated that Rb/E2F1 interacts (Altintas et al., 2012) and regulates *AR* expression, *AR* transcription and AR target gene expression (Gao et al., 2016; Sharma et al., 2010) to facilitate the development of hormone-refractory prostate cancer. Interestingly, in a *Drosophila* spinal and bulbar muscular atrophy model, it was observed that the *Drosophila* Rbf/E2F1 axis functionally interacts with human polyglutamine repeat expansion AR mutants to mediate neurodegeneration in *Drosophila* eyes (Nedelsky et al., 2010; Suzuki et al., 2009). Although it has not been possible to assess the highly cell type-specific physical interaction between EcR and Rbf/E2F1 because of the small number of SCs, it seems likely that part of Rbf/E2F1-dependent regulation of EcR and its hormone-independent signalling involves physical interaction between the EcR and these cell cycle regulator proteins. Importantly, while the control of SC nuclear growth by the BMP/EcR axis and cell cycle regulators appears complex, the regulation of endoreplication is absolutely dependent on all elements of this network and specifically involves hormone-independent EcR activity. Overall, the physiological control of SC endoreplication in mated male flies seems to share many regulatory parallels with the hormone-independent, AR-induced growth and proliferation observed in pathological CRPC, suggesting that SC endoreplication may be used as a model for investigating genetic interactors involved in CRPC.

## Materials and Methods

### *Drosophila* Stocks and fly husbandry

Following fly stocks were obtained from the Bloomington *Drosophila* Stock Center, unless other source is mentioned: esg^ts^F/O (*w*^−^; *esg-GAL4, UAS-GFP_nls_; act*>*CD2*>*GAL4, UAS-FLP*; gift from B. Edgar) (Jiang et al., 2009), *UAS-EcR-RNAi* (TRiP.JF02538) (Colombani, 2005), *UAS-EcR-C* (Cherbas, Hu, Zhimulev, Belyaeva, & Cherbas, 2003), *UAS-CycE-RNAi* (TRiP.GL00511), *UAS-CycE* (Richardson, O’Keefe, Marty, & Saint, 1995), *UAS-p53* (gift from J. Abrams), *UAS-Rbf^CA^* (*w[*\**]; P{w[*+*mC]*=*UAS-Rbf.280}3/TM3, Sb[1]*) (Xin et al., 2002), *UAS-Mud-RNAi* (TRiP.HMS01458) (Elkahlah, Rogow, Ahmed, & Clowney, 2020), *UAS-CycD/cdk4* (gift from B.Edgar) (Datar, Jacobs, De La Cruz, Lehner, & Edgar, 2000), *UAS-CycD-RNAi* (TRiP.HMS00059) (Kim, Jang, Yang, & Chung, 2017), *UAS-Rbf-RNAi* (TRiP.HMS03004) (Greenspan & Matunis, 2018), *UAS-E2F1/Dp*, *UAS-E2F1-RNAi* (TRiP. JF02718) (Zhang et al., 2017), *UAS-Tkv^act^* (gift from K. Basler) (Nellen, Burke, Struhl, & Basler, 1996), *UAS-Dad* (gift from D. Bennett) (Tsuneizumi et al., 1997), *UAS-shd-RNAi* (Leiblich et al., 2019). Flies were reared in standard cornmeal agar medium and experimental crosses were maintained either at 18°C or 25°C. For virgin experiments, newly eclosed virgin males of the required genotype were collected and kept on food with or without 0.2mM EdU (see below). For multiply mated experiments, each male was placed in individual vials with ten virgin *w^1118^* females (Leiblich et al., 2019, 2012; Redhai et al., 2016).

### Immunohistochemistry and microscopy

The protocol used for fixing and immunostaining the AG were performed as reported in our previous works (Corrigan et al., 2014; Leiblich et al., 2019, 2012; Redhai et al., 2016). Fixed samples were blocked with PBSTG (PBST with 10% Goat Serum) for 30 min at room temperature and were incubated overnight with primary antibody diluted in PBSTG at 4°C. They were washed in PBST for 6 × 10 min before being incubated with secondary antibody (1:400 in PBST) (ThermoFisher) for 30 min at room temperature. Primary antibodies Anti-FAS3 (Developmental Biology Hybridoma Bank (DSHB); 1:10) and anti-EcR (DSHB; 1:10) were used in conjunction with fluorescent Alexa-555- (ThermoFisher; 1:400) and Cy3- (Jackson Laboratories; 1:400) conjugated donkey anti-mouse secondary antibody respectively. The glands were imaged using an upright Zeiss LSM 880 Airy Scan confocal microscope. The Zeiss Plan-Apochromat 63X/1.4 NA Oil objective (Carl Zeiss) was used for 63X magnification and Zeiss Plan-Apochromat 40X/1.3 NA Oil objective (Carl Zeiss) was used for 40X magnification. Immersion oil of refractive index 1.514 (Cargill labs) was used.

### SC nuclear growth assay

Z-stack images of the whole SC (3 per gland) were acquired by confocal microscopy as discussed above. The sum of all stacked images was obtained and analysed using Fiji. For each selected SC, the sum of the maximum areas of both nuclei and of an MC adjacent to the SC was calculated. In the case of mononucleate cells (e.g. Rbf^CA^), the area of the single nucleus was calculated. SC Nuclear Area/MC Nuclear Area was used as a measure of SC growth (Leiblich et al., 2019, 2012).

### EdU incorporation assay

The Click-iT™ Plus EdU Cell Proliferation Kit for Imaging (Invitrogen; Alexa Fluor™ 594 dye) was used to detect DNA replication. EdU is a synthetic analogue of thymidine. Flies were maintained on medium prepared by mixing standard yeast-cornmeal agar medium with a 80mM EdU (Cambridge Bioscience) stock solution after at 60°C (diluted in PBS, as per the manufacturer’s instructions) to reach a final EdU concentration of 0.2mM; 750μL of the 80 mM EdU stock solution was used for every 300 mL of yeast-cornmeal agar. In order to detect EdU incorporation, flies fed on EdU-containing food were dissected, but the testes were kept intact as a positive control for EdU staining as the adult stem cell niches in the testes undergo DNA replication and incorporate EdU (de Cuevas & Matunis, 2011). Once suspended in PBST, tissues were transferred to PBSTG (and incubated for 45 minutes at RT to block, before being washed with PBST. DNA was labelled by adding the Click-iT^®^ reaction mix (prepared following manufacturer’s instructions) to the vials and left to incubate for 30 min at room temperature, away from light. Glands were washed in PBST for 3 × 10 min and then re-suspended in PBS. Only AGs from flies whose testes were positively stained for EdU staining were included for the analysis. The total number of SCs were quantified by counting the number of GFP-containing SCs (nGFP from esg^ts^F/O driver). Percentage of SCs which incorporated EdU was then used as a measure for endoreplication in the gland.

### Statistical analyses

Either the mean or geometric mean of SC:MC nucelar area or mean rank of % of EdU incorporated were compared across different genotypes. The normality or log-normality of the data were checked using the Shapiro-Wilk normality test and D’Augostino and Pearsons’ normality test. For log-normal data, further analyses were undertaken using log-transformed data. Bartlett’s homogeneity of variance test was used to check for similarity of variances (Equation 1); if variances were similar, One-way ANOVA followed by Tukey’s HSD post-hoc test was used and if variances were dissimilar, Welch ANOVA followed by Games-Howell post-hoc test was used. If the data were not either normally or log-normally distributed, Kruskal-Wallis test followed by Dunn’s post-hoc test was used. When the data are log-transformed, the analyses conducted compare the geometric means of the data rather than the arithmetic means.

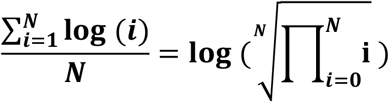

Where 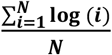 is the arithmetic mean of the log-transformed values and 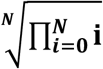 is the geometric mean of the actual data.

Therefore, when log-normal data are represented in a scatterplot graph, we show the geometric means and the corresponding geometric standard deviations.

## Acknowledgements

We are grateful to the Bloomington Drosophila Stock Center and Vienna Drosophila Stock Center for flies and to the Developmental Studies Hybridoma Bank (DSHB), Iowa, for EcR antibodies. Microscopy was undertaken in the Micron Oxford Advanced Bioimaging Unit.

## Notes

### Competing Interest Statement

The authors have declared no competing interest.

