## Supplementary Figures for "Rbf/E2F1 control growth and endoreplication via steroid-independent Ecdysone Receptor signalling in *Drosophila* prostate-like secondary cells"

**Supplementary figure legends:**

**Figure S1: Role of CycE and CycD in SC growth and endoreplication regulation**

(A-D) Images show distal tip of AGs from 6-day old adult males expressing nuclear GFP alone (control; this also stains the cytosol) or in combination with additional transgenes in SCs under the control of the $esg^{ts}$F/O driver and stained for EdU incorporation in SCs. Nuclei are stained with DAPI (blue). Red arrows point to $EdU^{+}$ SC nuclei and yellow arrows point to $EdU^{-}$ SC nuclei for each transgene. (E-F) Histograms depicting the mean ratio of the size of SC nuclei relative to neighbouring MC nuclei (E) and mean % of EdU+ SCs per gland (F) of virgin and mated flies expressing no other transgene or *CycE*-RNAi in SCs. Knocking down *CycE* completely inhibits SC endoreplication, but does not completely suppress SC growth that occurs after mating. Welch ANOVA; Games-Howell post hoc test (E). (G-J) Images show distal tip of AGs from 6-day old adult males expressing nuclear GFP alone (control; this also stains the cytosol) or in combination with additional transgenes in SCs under the control of the $esg^{ts}$F/O driver. (G'-J') Images show lower magnification of EdU stained AGs of control glands or glands expressing transgenes from 6-day old adult males. AGs are outlined in dotted white lines. Kruskal Wallis test; Dunn's post hoc test (F). n≥15 cells (E); n≥9 glands (F). Scale bars correspond to 20 µm (A-D, G-J) and 50 µm (G'-J'). The error bars show the standard deviation within the sample. 0.01<*p<0.05; 0.001<**p≤0.01; 0.0001<***p≤0.001; ****p≤0.0001.

**Figure S2: Role of Mud and E2F1 in the regulation of SC nucleation state**

(A-B) Images show distal tip of AGs from 6-day-old adult virgin males of glands expressing

*E2F1*-RNAi under the control of the esg$^{ts}$F/O driver (A) and glands expressing *E2F1*-RNAi under the control of the Acp26Aa-GAL4 driver (B) which activates nuclear GFP production (stains cytoplasm in A,C). (A-B) Dashed red ellipses mark the outline of mononucleate SCs; dashed yellow ellipses mark the outline of binucleate SCs. Scale bars correspond to 50 µm.

**Figure S3: Validation of SC nuclear growth assay for glands containing mononucleate and binucleate cells**

Histogram depicting the mean cellular area of SCs from control glands or glands expressing Rbf$^{CA}$ in SCs. The cellular area of SCs expressing Rbf$^{CA}$ is smaller than SCs from control glands, similar to the trend observed with the SC nuclear area assay (Fig.3 I). Mann-Whitney test; $n \geq 43$ cells; ****$p < 0.0001$.

**Figure S4: Role of E2F2 in SC growth and endoreplication regulation**

(A-D) Images show distal tip of AGs from 6-day old adult males expressing nuclear GFP alone (control; this also stains the cytosol) or in combination with additional transgenes in SCs under the control of the esg$^{ts}$F/O driver and stained for EdU incorporation in SCs. Nuclei are stained with DAPI (blue). Red arrows point to EdU$^{+}$ SC nuclei and yellow arrows point to EdU$^{-}$ SC nuclei for each transgene. (E-F) Histograms depicting the geometric mean ratio of the size of SC nuclei relative to neighbouring MC nuclei (E) and mean % of EdU+ SCs per gland (F) of virgin and mated flies expressing no other transgene or *E2F2*-RNAi in SCs. Knocking down *E2F2* does not affect SC growth or endoreplication in either virgin or mated males. One-way ANOVA on log-transformed data; Tukey’s HSD post-hoc test (E). Kruskal-Wallis test; Dunn’s post-hoc test (F). $n \geq 18$ (E); $n \geq 6$ (F). $0.0001 <$***$p \leq 0.001$.

**Figure S1:**

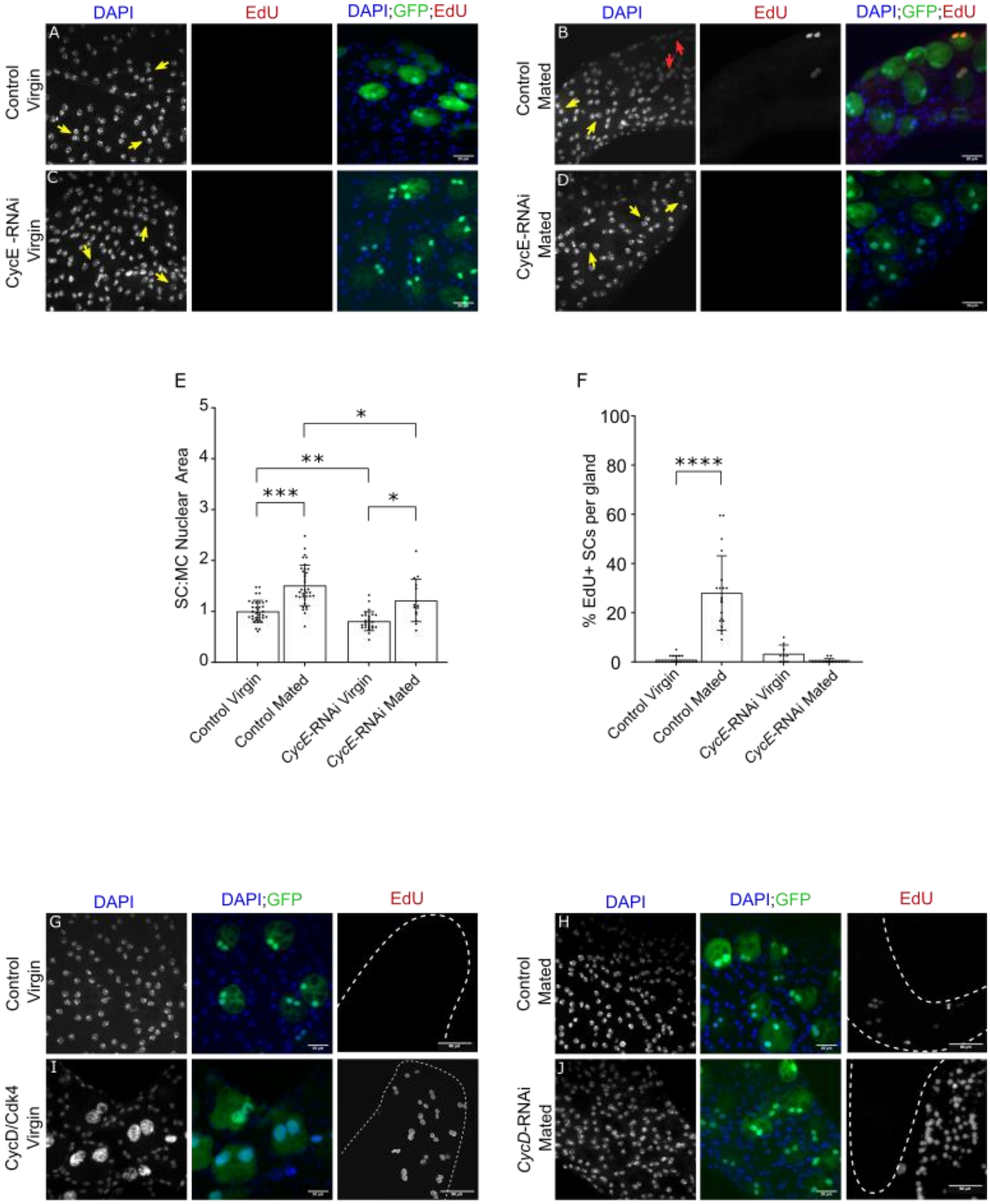

DAPI
EdU
DAPI;GFP;EdU
A
B
C
D
Control Virgin
Control Mated
CycE -RNAi Virgin
CycE-RNAi Mated
E
F
SC:MC Nuclear Area
% EdU+ SCs per gland
***
**
*
****
Control Virgin
Control Mated
CycE-RNAi Virgin
CycE-RNAi Mated
DAPI;GFP
G
H
I
J
CycD/Cdk4 Virgin
CycD-RNAi Mated

**Figure S2:**

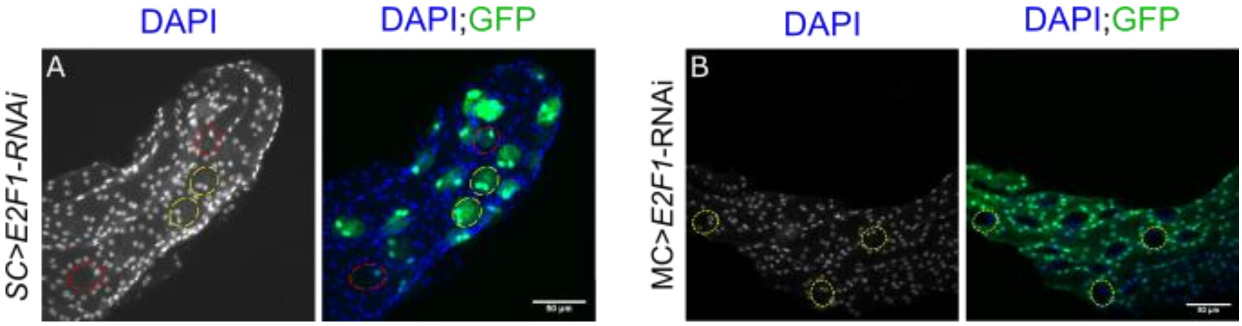


**Figure S3:**

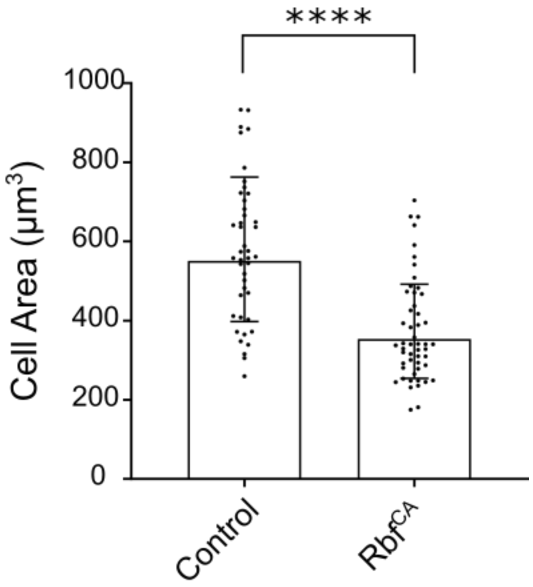

**Figure S4:**

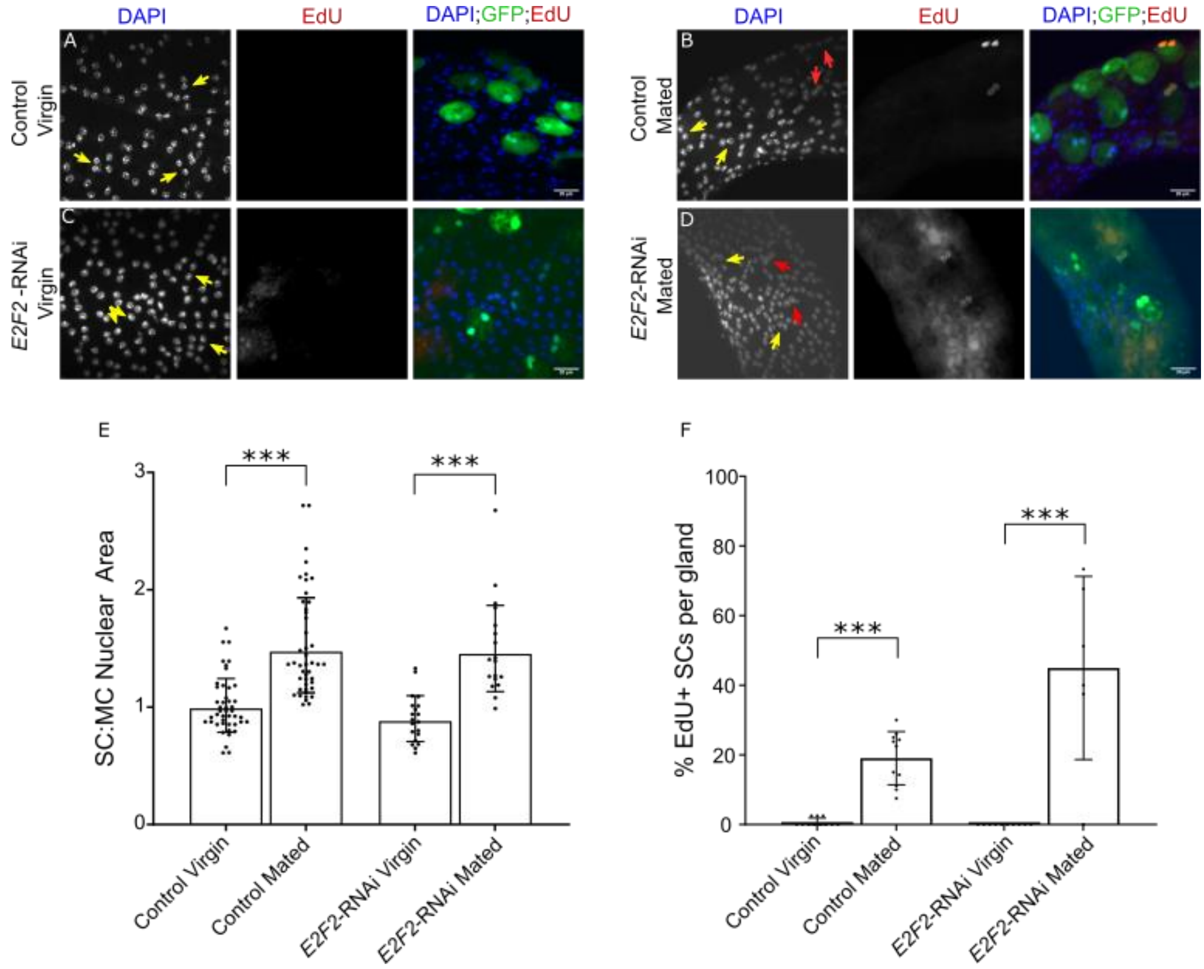
DAPI
EdU
DAPI;GFP;EdU
A
B
C
D
Control Virgin
Control Mated
E2F2 -RNAi Virgin
E2F2-RNAi Mated
E
F
SC:MC Nuclear Area
% EdU+ SCs per gland
***
Control Virgin
Control Mated
E2F2-RNAi Virgin
E2F2-RNAi Mated
